# A normative study of modified spatial context memory test in middle and older individuals

**DOI:** 10.1101/584136

**Authors:** Hsuan-Min Wang, Yo-Ping Huang, Hsun-Yu Kuo, Hung-Chou Kuo

**Author notes:** These authors contributed to this work equally (Hsuan-Min Wang, Yo-Ping Huang). Address correspondence and reprint requests to Hung-Chou Kuo, M.D., Department of Neurology, Chang Gung Memorial Hospital & Chang Gung University, 5 Fuxing St., Guishan Dist., Taoyuan, Taiwan 33333.

## Abstract

In view of the spatial context memory function of hippocampus complex region, we designed a modified spatial context memory test (SCMT) and try to early identify amnesic mild cognitive impairment.

All participants with non-dementia were recruited and divided 2 age groups (55-65 years and 66-75 years) and 3 education levels (6-9 years, 10-12 years and more than 12 years). The mini-mental state examination, visual associative memory test, visual construction retention test, and logical memory subtest of the Wechsler memory scale-III were used to evaluate the cognition state of the individuals. Spatial-context memory test version I with combination of a spatial memory paradigm and a real-life event included 3 subtests of navigation, scene-event association and people-object association.

A total of 147 individuals were confirmed to be normal in cognition in the assessment of the neuropsychological test battery. Regardless of age or level of education, there was no significance in perseveration errors and the retrieval of learned-well navigation information. In the Scene-Event Association test, the subjects with a low level of education seemed to have relative difficulty to quickly classify new information and establish the effective cues to retrieval. The subjects with a high level of education, the performance of spatial-context memory were negative correlation to the age. In the People-Object association test, the subjects in all six groups made no perseveration errors, but older subjects required more time to retrieval, and this situation was more prominent in the subjects with high level education.

We establish a normal age‐ and education‐adjusted SCMT score in the middle and elder individuals.

## Introduction

Persons with amnestic mild cognitive impairment (a-MCI) are generally believed to be at high risk for Alzheimer’s disease (AD) [1] and a-MCI is thought to represent recent memory impairment caused by the destruction of the neurons of the perirhinal and parahippocampus by neurofibrillary tangles [2,3]. According to Braak’s and Braak’s pathological process of Alzheimer’s disease (AD), a progression of neurofibrillary pathology from the transentorhinal and entorhinal cortex to the hippocampus, and then to the remaining limbic system before involving other cortical regions [2]. Previous functional magnetic resonance imaging (MRI) study has indicated that the perirhinal area processes item memory, especially in coding familiarity, the parahippocampus processes spatial-context information, and the hippocampus handles spatial navigation [4-6]. Previous functional MRI study has revealed that episodic encoding is mediated by distinct encoding processes, and suggests that perirhinal region mechanisms support the encoding of the individual elements of an episode, whereas the parahippocampus is involved in the processing of spatial relationships and in encoding of spatial context. The hippocampus supports domain-general relational processing and activation in spatial navigation [7]. According to neuropathological and functional imaging findings and our study [8] we hypothesized that impairment of spatial-context memory and spatial navigation are positively correlated with progressive degeneration of hippocampal structures; patients in very early neurodegenerative stages should have problems in the spatial-context memory tests. We therefore used computer software to establish a test of spatial-context memory, gathered normal data from a normal group, and hope to use the resulting data as a basis for comparing the test performance of persons with a-MCI and early AD, and trying correlations with imaging study.

## Materials and methods

In this study we based on the Microsoft visual studio 2017, Sql Server 2016 framework and Unity5 software to make a computer program call spatial-context memory test version I (SCMT-I).

### Subjects

We divided the subjects in this study into two groups with respective ages of 55-65 and 66-75 years, and three groups in accordance with levels of education (6-9 years of education, 9-12 years of education and over 12 years of education). This study recruited subjects between the ages of 55 and 75 years through posters, and excluded those who had existing neurodegenerative diseases, severe psychosis, severe visual impairment, and severe hearing impairment. The person with previous serious brain injuries, histories of epilepsy or cerebrovascular disease, and other systemic diseases that might affect cognitive function were also excluded. All subjects were informed in advance of the research procedures, research content, and then medical rights in detail, and written consent was obtained from all participants in accordance with protocol approved by Chang Gung Memorial Hospital (IRB: 201601546B0 and IRB: 201601546B0C601). This study recruited a total of 150 subjects, of whom 3 subjects dropped out of the test half-way due to lack of patience, and were consequently not included in statistical analysis. A final total of 147 subjects were therefore enrolled in this study.

### Procedures

All subjects in this study received a neuropsychological test battery, which included the Mini-Mental State Examination (MMSE), Visual Associative Memory Test (VAMT), Visual Construction Retention Test (VCRT), Logical Memory Test (LMT) of the Wechsler Memory Scale-III(WMS-III), and SCMT-I. Each subject was tested by a trained neuropsychologist in a standard testing environment, and the test took approximately 80 minutes to complete. Among the tests in the battery, MMSE, VAMT, VCRT, and LMT were administered to exclude the possibility that subjects had existing mild cognitive impairment or early dementia. Because this study sought to select subjects who were close to normal as much as possible for the purpose of analysis, after completing data collection from the 147-subject sample, we used the results from the neuropsychological test battery to establish three-stage selection criteria (Table 1). In this process, the first step was to use MMSE and VAMT results to select subjects for analysis in the second stage, where only subjects with an MMSE score of ≧26 and a total VAMT score of >9 entered the second stage. The second step consisted of selection of those subjects who had the scale score ≧8 in retention of logical memory test to enter the third stage. In the third step, only subjects with the scale score ≧8 in the first recall of logical memory scores were included in the final statistical analysis.

**Table 1.** Three-step of neuropsychological tests for normal short team memory function

| Total sample |  | 147/150 (3 subjects were drop out of this study) |  |
| --- | --- | --- | --- |
| Sampling inspection | Condition | Rule out | Sample |
| step 1 | VAMT total score > 9 | 25 | 122 |
| step 2 | Retention of LM < 8 (SS) | 8 | 114 |
| step 3 | LM1 < 8 (SS) | 6 | 108 |

VAMT, visual associative memory test; LMT, logical memory test; LMT1, first recall of logical memory test; SS, scale score of WMS-III;w stimulation/ total errors

### Spatial-context memory test version I

When administering the SCMT-I test, the subjects sat in front of a computer desk so that they could see all test stimuli as clearly as possible without interfering reflections. The test stimuli were uniformly displayed on the computer. The test including instructions and the subjects’ response time was about 30 min. The SCMT-I test (Figure 1) contains three parts: (1) Navigation test: while a stimulus was shown, the subjects were told that they must enter a city with a mission of withdrawing money from a bank. As they travelled through the city, the subjects encountered several corners near buildings at which they must turn. Arrows guided the subjects in the direction they must take, and the subjects were asked to remember the route. The test had a total of 6 corners. A question session began immediately after the stimulus was presented, at which time the subjects were asked to enter the same city and tried to remember the same route to the bank. The program automatically displays again if the subjects make the wrong choice of any one of the corners, and the test ended after the subjects answered two consecutive sets of 6 questions correctly. The test allowed a total of 10 trails, and the test was stopped if a subject cannot complete it correctly two consecutive sessions within 10 trails (Figure 1(a)).

**Figure 1.**
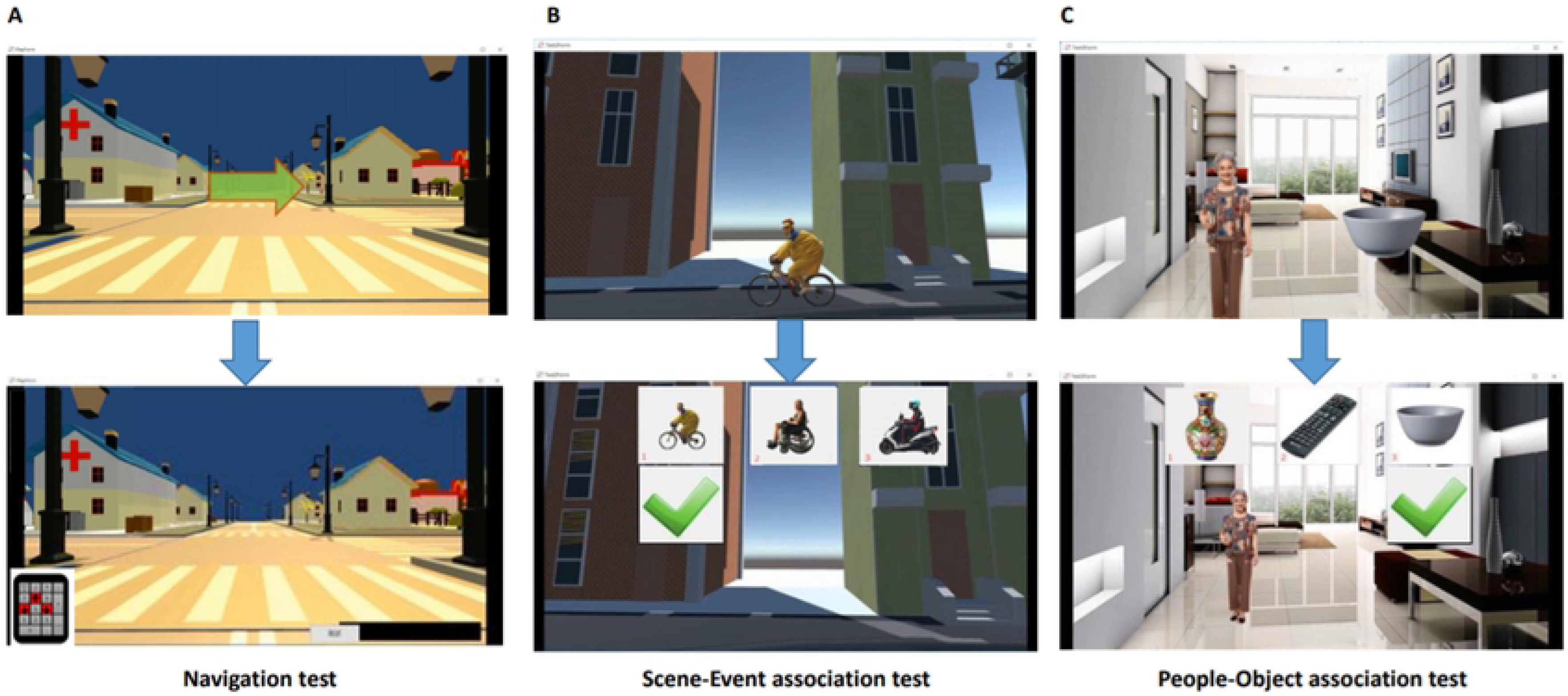
Sampling three subtests of spatial-context memory test version I including (a) navigation, (b) scene-event association, and (c) people-object association.

(2) Scene-event association test: While a stimulus was presented, the subjects were told that they must enter another city, where they would see a specific event at different locations as they proceeded. In this process, the subjects were asked to pay attention and remember what they saw. A question period began immediately after the end of the stimulus. During this period, the subjects were told to enter the same city again, where three possible event options were presented at a specific location; of these three options, one actually presented at this location, another belonging to the other scene not associated with this location, and the other is a new stimulation which was never presented before. The subjects were asked to try to recall, and use the keyboard to make their choice. If a subject made a wrong answer, the program displayed an error message and told the subject what the correct answer was. This test had a total of 12 questions; if they made any errors, the program will automatically display again. The subjects were asked to try again until all 12 questions have been answered correctly in two consecutive sessions. The subjects had 6 chances to complete the test; if a subject cannot complete the test successfully in two consecutive times after 6 trails, the test was stopped (Figure 1(b)).

(3) People-object association test: While a stimulus was presented, the subjects were told that they must enter another city, where they entered different buildings and met different people, and then obtained certain object from this people. The subjects were asked to pay attention and remember what they saw. A question period began immediately after the end of the stimulus. During this period, the subjects were told to enter the same city and building again, where they would see the people that they had met before and then three object options presented. They were asked to make choice which one they had reached from this people; of the three options, one actually presented from this people, another associated with the other people in another building not reached from this people, and the other is a new stimulation which was never presented before. The subjects were asked to recall and use the keyboard to make a choice. If a subject made a wrong answer, the program displayed an error message and told the subject the correct answer. This test had a total of 10 questions; if they made any errors, the program will automatically display again. The subjects were asked to try again until all 10 questions have been answered correctly in two consecutive sessions. The subjects had 6 chances to complete the test; if a subject cannot complete the test successfully in two consecutive sessions after 6 trails, the test was stopped (Figure 1(c)).

### Data collection

The subjects’ each response of SCMT-I will be recorded by program. In this study, answering all the questions correctly in two consecutive sessions indicated that the subjects were learned-well. In the (1) Navigation test: (i) the program separately recorded the total response time which the subjects answered all 6 questions correctly in two consecutive sessions; (ii) the subjects’ response time at each corner; (iii) the program will record each question answered being correct or wrong. Every wrong response was classified as either non-perseveration errors or perseveration errors; and (iv) the program also recorded how many trails the subjects took to learn well.

(2) Scene-event association test: (i) the program separately recorded the total response time which the subjects answered all 12 questions correctly in two consecutive sessions; (ii) the program would record each question response time from options appearance to subject making choice; (iii) the program would record each question answered being correct or wrong. The program also classified erroneous responses into the following four error types: (a) a non-perseveration error from other scene not associated with this event; (b) non-perseveration new stimuli error; (c) a perseveration error from other scene not associated with this event; and (d) perseveration new stimuli error; and (iv) the program also recorded how many trails the subjects took to learn well.

(3) People-object association test: (i) the program separately recorded the total response time which the subjects answered all 10 questions correctly in two consecutive sessions; (ii) the program would record each question responses time from options appearance to subject making choice; (iii) the program would record each question answered being correct or wrong. The program also classified erroneous responses into the following four error types: (a) a non-perseveration error from other scene not associated with this event; (b) non-perseveration new stimuli error; (c) a perseveration error from other scene not associated with this event; and (d) perseveration new stimuli error, and (iv) the program also recorded how many trails the subjects took to learn well.

### Statistical analysis

Differences in patient demographics and neuropsychological test results between the groups were determined using one-way ANOVA. All statistic assessments were two sided and P ≤ 0.05 was considered statistically significant. All statistical analysis was performed using SPSS 15.0 statistical software (SPSS Inc., Chicago, IL).

## Results

The results of the neuropsychological test battery are shown in Table 2. Table 3 demonstrates the navigation subtest results of SCMT-I that 2 subjects in the 55-65-years-of-age group failed to answer correctly all questions in 10 trails that resulted in a failure rate of 3.28%. In the 66-75-years-of-age group, only one subject failed to answer correctly all questions after 10 trails, and the failure rate is 2.13%. The subjects in all six groups the secondary total complete time was faster than first. The difference in total complete time between the first and second session seemed to expand associated with education level, but post hoc comparison revealed that no significant differences between any level of education and any age group. In addition, the results further reveal new information that the subject could remember in the first learning (first responses correct) was reduction with age (t:2.62, 95% C.I.: 0.24-1.77). The test results also reveal the perseveration errors were rare presentation either in difference education level or different age group.

**Table 2.** Individual demographic and neuropsychological test battery

| Groups: Age /Education (years) | 55-65/6-9 | 55-65/9-12 | 55-65/12↑ | 66-75/6-9 | 66-75/9-12 | 66-75/12↑ |
| --- | --- | --- | --- | --- | --- | --- |
| Number of subjects | 21 | 22 | 18 | 18 | 12 | 17 |
| Sex,% Male: Female | 29:71 | 59:41 | 56:44 | 39:61 | 42:58 | 29:71 |
| MMSE (SD) [30] | 27.57(1.43) | 28.5(0.91) | 28.94(1.00) | 27.94(1.43) | 28.58(1.16) | 28.65(1.46) |
| VAMT1 (SD) [6] | 5.48(0.75) | 5.5(0.67) | 5.72(0.46) | 5.56(0.70) | 5.50(0.52) | 5.06(0.75) |
| VAMT2 (SD) [6] | 6.00(0.00) | 6.00(0.00) | 6.00(0.00) | 6.00(0.00) | 5.92(0.29) | 5.94(0.24) |
| TVAMT (SD) [12] | 11.52(0.68) | 11.5(0.67) | 11.72(0.46) | 11.56(0.70) | 11.42(0.67) | 11.00(0.79) |
| VCRT (%) (SD) [100%] | 84.23(16.54) | 84.68(16.38) | 90.30(13.83) | 82.98(17.31) | 89(12.26) | 88.23(11.03) |
| LM1 (SD) [25] | 10.29(3.24) | 12.31(3.43) | 12.72(3.80) | 12.06(3.00) | 13.58(3.18) | 12.06(3.61) |
| LM2 (SD) [25] | 9.71(3.54) | 11.95(3.06) | 11.78(3.62) | 10.78(3.10) | 12.67(3.68) | 10.47(3.62) |
| Retention of LM (%) (SD) [100%] | 91.32(8.89) | 93.88(8.77) | 91.37(11.02) | 86.89(13.03) | 88.81(11.54) | 84.08(12.33) |

MMSE: Mini mental status examination; VAMT: Visual association memory test; TVAMT: Total visual association memory test; VCRT: Visual construction retention test; LM: Logical memory

**Table 3.** Navigation test

| Groups: Age /Education (years) | 55-65/6-9 | 55-65/9-12 | 55-65/12↑ | 66-75/6-9 | 66-75/9-12 | 66-75/12↑ |
| --- | --- | --- | --- | --- | --- | --- |
| Number of subjects | 21 | 22 | 18 | 18 | 12 | 17 |
| Failure (%) | 1 (4.76%) | 1 (4.55%) | 0 | 0 | 1 (8.33%) | 0 |
| 1<sup>st</sup> full completed time (sec) (SD) | 11.37 (6.25) | 11.20 (7.32) | 14.36 (12.10) | 13.18 (7.00) | 10.16 (3.80) | 10.68 (7.65) |
| 2<sup>nd</sup> full completed time (sec) (SD) | 8.20 (5.18) | 8.75 (10.39) | 9.37 (7.61) | 12.37 (10.57) | 8.9 (4.09) | 9.36 (4.43) |
| 2<sup>nd</sup>-1<sup>st</sup> difference (sec) (SD) | -2.82 (7.82) | -2.73 (8.06) | -4.99 (6.51) | -0.81 (13.58) | -1.23 (3.40) | -1.32 (5.70) |
| First responses (SD) [6] | 5.16 (1.61) | 5.40 (1.57) | 4.83 (1.76) | 4.35 (2.00) | 4.18 (2.27) | 3.88 (2.29) |
| Total errors (SD) | 1.11 (1.15) | 0.55 (1.05) | 0.72 (1.07) | 1.18 (1.19) | 1.00 (1.48) | 1.18 (1.19) |
| Perseveration errors (SD) | 0.05 (0.23) | 0.05 (0.22) | 0.22 (0.73) | 0.18 (0.39) | 0.27 (0.65) | 0.06 (0.24) |
| Perseveration errors (% , SD) | 1.75 (7.65) | 1.25 (5.59) | 6.94 (20.66) | 7.84 (17.79) | 8.18 (18.34) | 2.94 (12.13) |
| Learning times (SD) | 2.31 (1.38) | 1.65 (1.18) | 1.78 (1.11) | 2.19 (1.28) | 2.00 (1.48) | 2.29 (1.61) |

Percentage of Perseveration errors: Perseveration errors/ Total errors

The results of scene-event association test (Table 4) reveals that three subjects in the 55-65-years-of-age group failed to answer correctly all questions in 6 trails, and the failure rate is 4.92%. In the 66-75-years-of-age group, five subjects failed to answer correctly all questions after 6 trails, and the failure rate is 10.64%. The subjects in all six groups the secondary total complete time was faster than first, the post hoc comparison difference age group and education level revealed that there is no significant differences in time difference of completion. The results further indicated that, in the group with over 12 years of education, there was a significant difference in error rate between the 55-65 and 66-75-years-of-age groups (t:−2.60, 95% C.I.: −3.08- −0.37). In addition, further analysis of different types of errors revealed that the interference of new stimuli with memories increased with age in the group with 6-8 years of education. In the 55-65-years-of-age group, subjects with more than 12 years of education had a lower error rate (1.28 (0.57), 95% C.I.: 0.14-2.41), while further analysis of error type revealed the subject with 6-8 years of education was easily confused by the stimulation from other scene. The results of people-object association test (Table 5) indicate that no subjects in either the 55-65 or 66-75-years-of-age group failed in all 6 trails. The subjects in all six groups the secondary total complete time was faster than first. While the subjects in the 55-65-years-of-age group were able to complete the test faster in the second time than subjects in the 66-75 group. In the group with 6-8 years of education, subjects in the 66-75-years-of-age group spent more time completing the test both the first and second session than those in the 55-65-years-of-age group. This result seemed to indicate that age is an important factor affecting the subjects’ ability to recall information associated with persons. However, further analysis indicated that subjects in the 55-65-years-of-age group compared to 66-75-years-of-age group had many trails to complete the tasks. This result may indicate that the significant difference in the completion time may not be attributable to age, but may instead be attributable to the practice effect from multiple instances of learning. In the group with over 12 years of education, compared with the 55-65-years-of-age group and 66-75-years-of-age group, the former spent significant less time to complete the second sessions. This result may indicate that age is an important factor in the ability to recall person-related information among subjects with a high level of education, and suggests that subjects with an older age may need to spend more time to recall well-learned information associated with persons.

**Table 4.** Scene-event association test

| Groups: Age /Education (years) | 55-65/6-9 | 55-65/9-12 | 55-65/12↑ | 66-75/6-9 | 66-75/9-12 | 66-75/12↑ |
| --- | --- | --- | --- | --- | --- | --- |
| Number of subject | 21 | 22 | 18 | 18 | 12 | 17 |
| Failure (%) | 3 (14.28%) | 0 | 0 | 3 (16.67%) | 0 | 2 (11.76%) |
| 1<sup>st</sup> full completed time (sec) (SD) | 46.85 (14.77) | 49.70 (21.53) | 54.73 (29.20) | 56.76 (18.20) | 54.46 (27.83) | 48.96 (13.03) |
| 2<sup>nd</sup> full completed time (sec) (SD) | 46.71 (13.15) | 43.86 (15.70) | 48.26 (23.09) | 50.53 (11.32) | 42.63 (13.85) | 45.45 (14.41) |
| 2<sup>nd</sup>-1<sup>st</sup> difference (sec) (SD) | -1.20 (7.01) | -5.85 (10.82) | -6.47 (11.52) | -7.93 (15.15) | -12.76 (17.92) | -3.50 (7.10) |
| First correct responses (SD) [12] | 10.61 (1.24) | 10.57 (1.29) | 11.05 (0.94) | 10.00 (1.41) | 10.58 (1.78) | 10.53 (1.68) |
| Total errors (SD) | 2.89 (1.32) | 2.43 (2.16) | 1.61(1.38) | 4.73 (4.00) | 3.17 (2.62) | 3.33 (2.38) |
| Errors from other scene (SD) | 1.89 (1.13) | 1.33 (1.29) | 0.83 (0.86) | 2.07 (1.94) | 2.25 (2.01) | 1.40 (0.91) |
| Errors from other scene (% , SD) | 70.46 (32.33) | 42.75 (36.68) | 42.22 (40.31) | 42.60 (35.60) | 50.32 (39.67) | 44.59 (34.48) |
| Errors from new stimulation (SD) | 0.89 (0.96) | 0.90 (0.83) | 0.72 (0.75) | 1.87 (1.50) | 0.42 (0.51) | 1.33 (1.29) |
| Errors from new stimulation (% , SD) | 27.03 (31.85) | 35.66 (35.52) | 34.44 (38.47) | 50.17 (36.07) | 15.63 (28.89) | 31.30 (29.94) |
| Perseveration of errors from other scene (SD) | 0 | 0 | 0 | 0.33 (0.72) | 0.33 (0.65) | 0.13 (0.35) |
| Perseveration of errors from other scene (% , SD) | 0 | 0 | 0 | 3.11 (6.48) | 5.71 (12.12) | 2.07 (5.73) |
| Perseveration of errors from new stimulation (SD) | 1.11 (0.32) | 0.19 (0.68) | 0.06 (0.24) | 0.47 (1.3) | 0.17 (0.39) | 0.47 (0.92) |
| Perseveration of errors from new stimulation (% , SD) | 2.50 (7.32) | 2.54 (8.29) | 1.11 (4.71) | 4.11 (9.74) | 3.33 (7.78) | 8.70 (16.02) |
| Learning times (SD) | 2.83 (1.20) | 2.62 (1.47) | 2.39 (1.14) | 3.07 (1.21) | 2.75 (1.54) | 2.87 (1.19) |

Percentage of errors from other scene: Errors from other scene/ total errors; Percentage of errors from new stimulation: Errors from new stimulation/ Total errors; Percentage of perseveration of errors from other scene: perseveration of errors from other scene/ total errors; Percentage of perseveration of errors from new stimulation: perseveration of errors from new stimulation/ total errors

**Table 5.** People-object association test

| Groups: Age /Education (years) | 55-65/6-9 | 55-65/9-12 | 55-65/12↑ | 66-75/6-9 | 66-75/9-12 | 66-75/12↑ |
| --- | --- | --- | --- | --- | --- | --- |
| Number of subject | 21 | 22 | 18 | 18 | 12 | 17 |
| Failure (%) | 0 | 0 | 0 | 0 | 0 | 0 |
| 1<sup>st</sup> full completed time (sec) (SD) | 22.85 (7.60) | 22.47 (8.06) | 23.89 (8.96) | 29.40 (10.47) | 24.97 (10.87) | 23.19 (5.02) |
| 2<sup>nd</sup> full completed time (sec) (SD) | 18.39 (5.46) | 17.30 (5.31) | 18.71 (7.02) | 23.88 (6.05) | 19.94 (6.15) | 21.95 (5.46) |
| 2<sup>nd</sup>-1<sup>st</sup> difference (sec) (SD) | -4.46 (4.61) | -5.16 (5.36) | -5.19 (4.27) | -5.52 (8.21) | -5.03 (5.80) | -1.24 (3.79) |
| First correct responses (SD) [10] | 9.86 (0.47) | 10.00 (0.00) | 10.00 (0.00) | 10.00 (0.00) | 9.75 (0.87) | 10.00 (0.00) |
| Total errors (SD) | 0.24 (0.54) | 0 | 0 | 0 | 0.25 (0.87) | 0.12 (0.33) |
| Errors from other scene (SD) | 0.19 (0.51) | 0 | 0 | 0 | 0.17 (0.58) | 0.12 (0.33) |
| Errors from other scene (% , SD) | 14.29 (35.86) | 0 | 0 | 0 | 5.56 (19.25) | 11.76 (33.21) |
| Errors from new stimulation (SD) | 0.05 (0.22) | 0 | 0 | 0 | 0.08 (0.29) | 0 |
| Errors from new stimulation (% , SD) | 4.76 (21.82) | 0 | 0 | 0 | 2.78 (9.62) | 0 |
| Perseveration of Errors from other scene (SD) | 0 | 0 | 0 | 0 | 0 | 0 |
| Perseveration of Errors from other scene (% , SD) | 0 | 0 | 0 | 0 | 0 | 0 |
| Perseveration of Errors from new stimulation (SD) | 0 | 0 | 0 | 0 | 0 | 0 |
| Perseveration of Errors from new stimulation (% , SD) | 0 | 0 | 0 | 0 | 0 | 0 |
| Learning times (SD) | 1.19 (0.40) | 1.00 (0.00) | 1.00 (0.00) | 1.00 (0.00) | 1.08 (0.29) | 1.24 (0.66) |

Percentage of errors from other scene: Errors from other scene/ total errors; Percentage of errors from new stimulation: Errors from new stimulation/ Total errors; Percentage of perseveration of errors from other scene: perseveration of errors from other scene/ total errors; Percentage of perseveration of errors from new stimulation: perseveration of errors from new stimulation/ total errors

## Discussion

In everyday life, memories are chiefly stored as episodic memories, and the event-related content of these memories uses the background in time and space to construct their context. The temporal and spatial memory allows us to also recall the place where we saw someone, what was spoken, and what things happened at that time. The episodic memories formed basis of the time and space background are also termed context memory [9] because these memories are constructed in the hippocampal formation, problems forming or recalling these memories may represent cognitive function obstacles in patients with a-MCI. Context memories can be further classified into temporal-context memories and spatial-context memories. This study focused on spatial-context memory, and discusses the neural mechanisms that form these memories. If a better understanding of neural mechanisms can be obtained, this will facilitate the early detection of neurodegenerative disease like a-MCI.

Morris et al. revealed that the hippocampus and peripheral structures bear responsibility for animals’ spatial memories and navigation abilities [10]. In view of Darwin’s theory of evolution, the reason spatial memory is so important is that the absence of these memories would cause severe problems for animals in the wild. Without these memories, animals might forget where they had obtained food or water, and would have to spend much more time searching for sources of food or water. An animal might forget where it encountered danger and repeatedly put itself in great jeopardy. Because of the importance of memory to animals in the wild, evolution gradually created “place cells” facilitating the recognition of places. Morris et al. also verified that place cells exist in the hippocampus, and are linked with the hippocampus’ surrounding structure [10].

O’Keefe and Burgess discovered that when mice walked to a place in their environment where they previously obtained a reward, their place cells were stimulated after information on environmental characteristics entered their brain, and the place cells then stimulated goal cells. This linkage enabled mice to remember what happened in that place, and allowed the location of goal cells below the place cells in the subicular complex or nucleus accumbens to be inferred [11]. Burgess et al. [5,6] suggested that the subicular complex should also contain the so-called “event cells,” which are linked with place cells in accordance with Hebb’s rule. When event cells are activated, they indirectly stimulate place cells, which enables the representation of the spatial context of past events [5,6]. Similarly, place cells can also indirectly stimulate event cells, which enable events associated with the place to be represented.

Functional imaging study has also revealed that the hippocampus and its peripheral structure plays an indispensable role when humans create and recall events associated with spatial context memories [5, 6, 9, 12, 13]. The results of the study also indicate that when humans create and recall memories associated with places, there is not only significant activity in the right side of the hippocampus, but such other areas as the left side of the hippocampus, parietal association cortex, frontal and posterior cingulate cortex, and prefrontal lobe also have significant activity. These phenomena can explain different aspects of episode of memories. For instance, the parahippocampus is not only involved in the recognition of spatial information concerning landmarks, linkage between this area and the hippocampus is also involved in memories concerning the locations of objects. In addition, previous study also discovered that the prefrontal lobe also has significant activity associated with episodic memories, which may be connected with recall strategies and the need to eliminate interference from similar situation [5,6]. As a consequence, research investigating spatial context can not only shed light on the importance of the hippocampus in animals’ spatial memory, but can also illuminate the role of the different hippocampus regions in the recall of episodic memories in humans.

With regard to the pathways of neurocognitive functions, and the process of transmission of visual information from the occipital lobe to the hippocampus through the posterior parietal cortex and ventral temporal cortex, two directional processes occur simultaneously. In the first process, spatial background information arrives at the parahippocampus via the posterior parietal cortex, and is ultimately stored after processing in the parahippocampus [14, 15]. In the second process, information concerning event and object characteristics is ultimately transmitted via the ventral temporal cortex to the perirhinal area for processing; linkage exists between the perirhinal area and parahippocampus chiefly to process the linkage between event and object characteristics and associated spatial background information [15]. After the processed information is sent to the hippocampus for classification and automatic linkage, it is transformed into event codes for storage. It can be inferred from these two directional processes that the parahippocampus mainly processes spatial background memories, while the perirhinal area chiefly processes memories concerning event and object characteristics, and checks whether there is a linkage between spatial background information and event or object characteristics.

Persons with aging commonly feel that learning new information is difficult, but this problem may be derived from other cognitive functions, such as attention, information processing speed, and working memory. However, such persons will encounter no major difficulties when using already learned knowledge or retrieving well-stored new information after a delay. The findings of our study support these arguments. While humans may be less likely to quickly recall information as they age, remote memories will not be easily forgotten, and this phenomenon is not affected by level of education. In addition, regardless of age and level of education, the subjects in this study made very few perseveration errors. This indicates that the subjects could remember previous errors, and could avoid repeating the same error on their next attempt. To summarize this study’s navigation test findings, regardless of age or level of education, persons with normal aging will face no significant obstacles to the delayed retrieval of well-learned navigation information. Furthermore, such persons will seldom make perseveration errors, explaining why the subjects in this study could remember their past errors, and avoid the repetition of similar errors. These results reflect the importance indication the normal aging people even in a new environment, they also can rely on learning to avoid getting lost.

The results on the scene-event association test indicated that in persons with normal aging and a relatively high level of education, age is a key factor affecting spatial-context memory. Many people complain subjectively of age-related memory loss. The results on this subtest also indicated the persons with low level of education may have an inherently sparing ability to use strategy for quickly classifying new information and establish effective cues to retrieval; therefore, when they get older, they might have limitations to immediately operate a lot of information that made them easier to be confused by similar situations.

The results of the people-object association test revealed that, in comparison with subjects with medium and higher levels of education, subjects in the 55-65 group with a low level of education (6-8 years) require relatively many trails of learning before they can fully remember information; these subjects also made relatively many errors, and these errors were chiefly caused by confusion with events in other scene. This indicates that, among subjects in the 55-65-yeares-of-age group, when subjects had a relatively low level of education, they readily confused spatial-context information, which may have been connected with their sparing ability to rapidly classify information and form representational cues facilitating retrieval. In the 66-75-yeares-of-age group, compared with subjects with high level of education, subjects with low level of education required more trials to remember new information. This diversity may be connected with the fact that the subjects with a higher level of education could better use various strategies to aid their memory. To summarize the results of the people-object association test, the older subjects spent more time to correctly recall information associated with persons, and this effect was most noticeable among the normal aging subjects with higher level of education. However, among subjects with a low level of education, the fact that the younger subjects could more quickly retrieve person-associated information may reflect a practice effect, and this phenomenon may also be due to more trail of learning had increased their retrieval speed.

It may be the persons with low level of education may have an inherently sparing ability to use strategy for quickly classify new information and establish effective cues to retrieval; therefore, when they get older, they might have limitations to immediately operate a lot of information, but if they can learn certain strategies to aid remember, it may facilitate the delayed retrieval.

As for neuropsychological assessment, there are still no highly effective instruments for distinguishing normal aging from early a-MCI. This study employed a computer program to establish a test of spatial-context memory, gathered normal values from a normal group, and then used the resulting data as a basis for comparing the test performance of persons with a-MCI and early AD with normal values, and determining correlations with imaging results.

## Acknowledgments

This study was sponsored by a grant (CORPG3G0061) from Chang Gung Memorial Hospital, Taipei, Taiwan and by a joint project between the National Taipei University of Technology and the Chang Gung Memorial Hospital under Grant NTUT-CGMH-106-05.

## Author Contributions

All the authors cooperated and contributed to the design and plan of the present study. H-C K and Y-P H was in charge of funding acquisition. H-M W, Y-P H and H-C K performed the project administration. H-Y K was a computer software designer. H-M W and H-C K was in charge of clinical evaluation. H-M W, H-Y K and H-C K was in charge of data curation. H-M W wrote original draft. H-C K were in charge of manuscript verifying.

